# Providing recombinant gonadotropin-based therapies that induce complete oogenesis and produce viable larvae from an immature teleost, flathead grey mullet (*Mugil cephalus*)

**DOI:** 10.1101/2020.06.04.132175

**Authors:** Sandra Ramos-Júdez, François Chauvigné, Wendy Ángela González-López, Hanna Rosenfeld, Joan Cerdà, Ignacio Giménez, Neil Duncan

**Affiliations:** IRTA, Sant Carles de la Ràpita Ctra. de Poble Nou km. 5.5, 43540 Sant Carles de la Ràpita, Tarragona, Spain; IRTA-Institute of Biotechnology and Biomedicine (IBB), Universitat Autònoma de Barcelona, Parc de Recerca UAB, Mòdul B, E-08193 Bellaterra, Barcelona, Spain; Israel Oceanographic Limnological Research, National Center for Mariculture, Eilat, Israel; Rara Avis Biotec, S. L., Valencia, Spain

**Keywords:** *Mugil cephalus*, flathead grey mullet, gametogenesis induction, eggs, larvae, rFsh, rLh

## Abstract

Under intensive captive conditions, wild-caught flathead grey mullet (Mugil cephalus) females remained arrested at early gonad development and no sperm could be obtained from males. With the aim to induce and complete oogenesis, induce the release of sperm and obtain fertilized eggs, adult female and male grey mullet were treated with M. cephalus single-chain recombinant gonadotropins (rGths), follicle-stimulating (rFsh) and luteinizing (rLh) hormones. In Experiment 1, fish were treated with a weekly dose of rFsh (15 μg kg−1), which in females significantly (P < 0.001) increased plasma concentration of 17β-estradiol and induced vitellogenic oocyte growth up to a maximum mean diameter of 425 ± 19 μm after 9 weeks of treatment. In Experiment 2, fish were treated with weekly injections of both rFsh and rLh at different doses (from 2.5 to 12 μg kg−1). Oocyte diameter reached 609 ± 5 μm, from which final oocyte maturation and ovulation was induced with 30 μg kg−1 of rLh and 40 mg kg−1 of progesterone. Good quality sperm (> 75% motile spermatozoa) was obtained from males in both experiments, and in Exp. 2 the addition of rLh induced the production of higher quantities of sperm that were used to fertilise the eggs. Although fertilisation was low (0.4 %), these fertilized eggs with embryo development produced viable larvae (71% hatching rate). This is the first report in a teleost species, to obtain larvae from eggs that were from immature females induced through to maturation with rGths. In comparison, control females remained arrested as immature fish and control males did not produce sperm. The study demonstrated that both rGths are effective to induce the entire process of oogenesis in sexually immature female grey mullet and to obtain flowing sperm from males, adding more data to confirm the roles of the Gths in teleost gametogenesis. This advance provides the bases of a therapy for the use in the aquaculture of teleost of commercial interest or the conservation of endangered species.

## 1. Introduction

The flathead grey mullet (Mugil cephalus) is a catadromous teleost with a worldwide distribution (between latitudes 40° North and South) (McDonough et al., 2005) that has been cultured for several centuries principally in some Asian countries and around the Mediterranean basin. Many positive attributes of grey mullet culture have made this species a suitable option for aquaculture. Grey mullet has fast growth (0.75 - 1 kg per year) (FAO, 2019), does not require dietary fish meal and oil and can be reared in a wide range of salinities and culture systems including polyculture (González-Castro and Minos, 2016). In addition, the final product marketed in various forms has good texture, taste (Yousif et al., 2010) and is an excellent source of omega-3 essential fatty acids (Khemis et al., 2019).

Although this species has a long history of culture, it exhibits different degrees of reproductive dysfunctions in both genders under captive conditions. These dysfunctions have limited the possibility to close the life cycle and, thus, culture has been based on the capture of wild juveniles (Yousif et al., 2010) or induced spawning of wild breeders (Abraham et al., 1999; Das et al., 2014; El-Gharabawy and Assem, 2006; Karim et al., 2016; Vazirzadeh and Ezhdehakoshpour, 2014). Two types of reproductive dysfunction have been described and can be placed in two categories; arrest in late or early stages of gametogenesis. Arrest in late stages of gametogenesis (maturation and ovulation) has been observed in recently caught wild mullet or wild mullet that were acclimated to ponds or large tanks (El-Greisy and Shaheen, 2007; Kuo et al., 1973; Yousif et al., 2010). This is the most commonly observed dysfunction in fish and can be controlled by hormonally inducing spawning (Mañanós et al., 2009; Zohar and Mylonas, 2001) as has been achieved for flathead grey mullet (Abraham et al., 1999; Das et al., 2014; El-Gharabawy and Assem, 2006; Karim et al., 2016; Vazirzadeh and Ezhdehakoshpour, 2014; Yousif et al., 2010). However, the spawning induction of these wild fish arrested in the late stages of gametogenesis does not represent a sustainable solution for mullet culture. The more severe reproductive dysfunction when development is arrested in the early stages of gametogenesis has been observed in wild and hatchery reared fish held in intensive culture conditions in the Mediterranean region. Females did not initiate vitellogenesis; remained at the primary growth stage or cortical alveoli stage (present study), or were arrested at early stages of vitellogenesis (Aizen et al., 2005). Males failed to initiate spermiation (De Monbrison et al., 1997; Yashouv, 1969) or produced highly viscous milt that could not fertilize the eggs (Shehadeh et al., 1973). These reproductive dysfunctions may be related to alterations in the endocrine control in the brain-pituitary-gonadal (BPG) axis.

In vertebrates, the pituitary gonadotropins (Gths), the follicle-stimulating hormone (Fsh) and luteinizing hormone (Lh), are generally accepted to be the central components of the BPG axis in the control of gonad development. Current knowledge in teleost suggest that the major role of Fsh is to promote gametogenesis from early stages through to late stages (vitellogenesis in females and spermatogenesis in males), while Lh is involved in gamete final maturation and release (ovulation and spermiation, in females and males, respectively) (Lubzens et al., 2010; Mañanós et al., 2009). The mechanism underlying the reproductive dysfunctions in Mediterranean captive mullets has been described as an inhibition by dopamine (DA) on the action of gonadotropin releasing hormone (GnRH) to release Gths in both females (Aizen et al., 2005) and males (Glubokov et al., 1994). Therefore, methods based on the mechanisms controlling gametogenesis are required to induce complete gonadal development, from early stages through to the late stages. In the case of males, 17α-methyltestosterone (MT) implants enhanced spermatogenesis and spermiation (Aizen et al., 2005; De Monbrison et al., 1997). In females, treatment with GnRH agonist (GnRHa) in combination with a DA antagonist (Aizen et al., 2005) or a single injection of recombinant Fsh produced in the yeast Pichia pastoris (Meiri-Ashkenazi et al., 2018) increased the number of vitellogenic females by promoting the release of Gths from the pituitary. However, hormonal therapies to enhance endogenous Lh release have been observed to be less effective when the pituitary Lh content was low (Yaron et al., 2009), indicating that alternative therapies may be required in these situations.

An alternative strategy to control gametogenesis in mullet as in other teleost, which would not require the availability of endogenous Gths from the pituitary, is the long-term use of recombinant Fsh and Lh (rFsh and rLh, respectively). This approach is nowadays possible through the production of large amounts of species-specific rGths in heterologous expression systems, such as the Drosophila S2 cell line (Kazeto et al., 2008; Zmora et al., 2007), the yeast Pichia pastoris (Aizen et al., 2007; Chen et al., 2012; Kamei et al., 2003; Kasuto and Levavi-Sivan, 2005; Palma et al., 2018; Sanchís-Benlloch et al., 2017), baculovirus silkworm larvae (Cui et al., 2007; Glubokov et al., 1994; Ko et al., 2007; Kobayashi et al., 2010, 2003; Meri et al., 2000), HEK293 cells (Kazeto et al., 2019) and Chinese hamster ovary (CHO) cells (Chauvigné et al., 2017; Choi et al., 2005; Giménez et al., 2015; Molés et al., 2011; Peñaranda et al., 2018; So et al., 2005). The application of rGths based therapies has shown promise to control gametogenesis in different teleost (Chauvigné et al., 2018, 2017; Giménez et al., 2015; Kamei et al., 2006; Peñaranda et al., 2018) and, therefore, could be an effective method to induce gametogenesis in cultured mullet arrested in the early stages of sexual maturation.

The present study aimed to use homologous single-chain rGths produced in CHO cells as the basis of a long-term hormone therapy to obtain viable offspring from sexually immature flathead grey mullet females and males that did not have flowing sperm.

## 2. Material and methods

### 2.1. Study animals and maintenance

Flathead grey mullet were used in two experiments to examine the effect of rGth hormone therapies. Experiment 1 examined the long-term effect of rFsh on vitellogenesis and hormone therapies (GnRHa+MET or hCG) previously used in grey mullet to induce final oocyte maturation and ovulation. Experiment 2 examined the effect of a combined rFsh and rLh therapy. In order to obtain sperm, males were administered rFsh (Exp. 1) or rFsh in combination with rLh (Exp. 2). Experiment 1 used wild-caught mullet from the Ebro River reared in 10-m^3^ tanks for 7-9 months in IRTA facilities (Sant Carles de la Ràpita, Spain). In Experiment 2, the broodstock was formed with wild-caught individuals reared for 19-21 months and individuals reared for 3 months in IRTA that were obtained from semi-extensive pond culture in the fish farm Finca Veta La Palma (Isla Mayor, Spain). All fish were tagged intramuscularly with a Passive Integrated Transponder (PIT) tag (Trovan®, ZEUS Euroinversiones S.L. Madrid, Spain) for individual identification. To determine the sex of individuals, a sample of gonadal biopsy was obtained through slight suction with a plastic catheter (1.67 × 500 mm; Izasa Hospital, Barcelona) inserted approximately 5 cm through the gonopore. Individuals were assigned as males if no oocytes were observed in the biopsies

One month before each trial, individuals were transferred to a 10-m^3^ tank in a recirculating system (IRTAmar®) under natural conditions and were gradually acclimatized from fresh water to sea water at 36 ‰ to provide the conditions for gonad development, as Tamaru et al. (1994) concluded that the rate of oocyte growth was lower in females maturing in fresh water. To evaluate in vivo dose-response of rFsh, fish were held for 21 days in May when temperature was controlled to 24 ± 1 °C and photoperiod was natural. During Experiment 1, completed from August to November, water temperature was controlled at 24 ± 1 °C. Photoperiod was ambient until October, when a constant 11L:13D (light:dark) was maintained through to the end of the experiment to avoid large changes of decreasing day length. Individuals were daily fed on a soft mixture of 20 % sardines, 20% hake, 15 % mussels, 10 % squid, 10 % shrimp, spirulina and 25 % a commercial broodstock diet (Mar Vitalis Repro, Skretting, Spain). In Experiment 2, completed from the end of July to mid-October, water temperature was also controlled at 24 ± 1 °C while photoperiod was ambient. Fish were fed a commercial marine fish broodstock diet (Brood Feed Lean, Sparos, Portugal) during five days a week at a daily rate of 1.5% of the body weight and two days a week with mussels and polychaetes. Prior to the experiments, fish had the same feeding regimens and were held in natural conditions of photoperiod and temperature.

The present experimental study has been approved by IRTA’s Ethics Committee for Animal Experimentation and the Animal Experimentation Commission from the Local Government (Dpt. de Territori i Sostenibilitat from the Generalitat de Catalunya). The study was conducted in accordance with the European Union, Spanish and Catalan legislation for experimental animal protection (European Directive 2010/63/EU of 22nd September on the protection of animals used for scientific purposes; Spanish Royal Decree 53/2013 of February 1st on the protection of animals used for experimentation or other scientific purposes; Boletín Oficial del Estado (BOE), 2013; Catalan Law 5/1995 of June 21th, for protection of animals used for experimentation or other scientific purposes and Catalan Decree 214/1997 of July 30th for the regulation of the use of animals for the experimentation or other scientific purposes). During all experimental procedures, for hormone administration and sampling, fish were first anaesthetised with 73 mg L-1 of MS-222 and placed in a tank with 65 mg L-1 of MS-222 for manipulation.

### 2.2. Cloning of M. cephalus Gths β and α subunits for rGths production

The pituitary gland was removed from sacrificed fish, frozen in liquid nitrogen, and stored at −80°C. Total RNA was purified using the GenEluteTM mammalian total RNA miniprep kit (Sigma-Aldrich) according to the manufacturer’s instructions, and cDNA synthesis was performed with 1 μg of total RNA following the manufacturer’s instructions of the 3’ RACE kit (Invitrogen). Polymerase chain reaction (PCR) was carried out as indicated in the 3’ RACE kit using partially degenerated forward primers for the Fshβ or α subunits, the common abridged universal amplification primer (AUAP) as reverse primer, and the EasyATM high-fidelity PCR cloning enzyme (Agilent Technologies, Santa Clara, CA, USA). The forward primer for each gene covered the translation initiation codon ATG and was designed based on sequences available in the GenBank repository for Epinephelus coioides (AY186242), Oreochromis niloticus (AY294015), Dicentrarchus labrax (AF543314), Acanthopagrus schlegelii (AY921613), Maylandia zebra (XM_004558042), Fundulus heteroclitus (M87014), Oryzias latipes (AB541981), Sparus aurata (AF300425), Amphiprion melanopus (EU908056), Chrysiptera parasema (KM509061), and Kryptolebias marmoratus (EU867505). For Fshβ, the forward primer was 5’-ATGCAGCTGGTTGTCATGGYAGC-3’, whereas for the α subunit the primer was 5’-ATGGGCTCMNTGAAAYCHVCTG-3. The Lh□ subunit was cloned using a degenerate forward primer covering the central region of the RNA (5’-CAAYCAGACRRTDTCTCTRGA), designed based on teleost sequences publically available (E. coioides, AY186243; Oreochromis niloticus, AY294016; Dicentrarchus labrax, AF543315; Acanthopagrus schlegelii, EF605276; Maylandia zebra, XM_004553532; Pundamilia nyererei XM_005741532; Fundulus heteroclitus, M87015; Cyprinodon variegatus, XM_015404196; Oryzias latipes, AB541982; Kryptolebias marmoratus, XM_017431834; Poecilia reticulata XM_008429103; Nothobranchius furzeri, XM_015975766; Xiphophorus maculatus, XM_005816155), and the reverse AUAP primer. The 5’ end of the cDNA was further amplified using RACE (5’ RACE kit, Invitrogen) and specific primers. In all cases, the PCR products were cloned into the pGEM-T Easy vector (Promega Biosciences, LLC, San Luis Obispo, CA, USA) and sequenced by BigDye Terminator Version 3.1 cycle sequencing on ABI PRISM 377 DNA Analyser (Applied Biosystems, Life Technologies, Carlsbad, CA, USA). The nucleotide sequence corresponding to the full-length Lh□, Fshβ and α subunit cDNAs were deposited in GenBank with accession numbers MF574169, MF574168 and MF574167, respectively. Single chain recombinant M. cephalus rFsh and rLh were produced by Rara Avis Biotec S.L. (Valencia, Spain) using in-house technology. Briefly, CHO cells where transfected with expression constructs encoding fusion proteins containing the entire coding sequence of M. Cephalus Fshβ (GenBank accession n° MF574168) or Lhβ (GenBank accession n° MF574169) subunit, the 28 carboxyl-terminal amino acids of the hCG β subunit as a linker, and the mature sequence of the M. cephalus glycoprotein hormone α subunit (GenBank accession n° MF574167). The secreted recombinant hormones were subsequently purified from the culture medium by ion exchange chromatography, concentrated and stored at −80°C until use.Flathead grey mullet were used in two experiments to examine the effect of rGth hormone therapies. Experiment 1 examined the long-term effect of rFsh on gametogenesis and was followed by Experiment 2 that examined the effect of a combined rFsh and rLh therapy. Experiment 1, used wild-caught mullet from the Ebro River reared in 10-m^3^ tanks for 7-9 months in IRTA facilities (Sant Carles de la Ràpita, Spain). In Experiment 2, the broodstock was formed with wild-caught individuals reared for 19-21 months and individuals reared for 3 months in IRTA that were obtained from semi-extensive pond culture in the fish farm Finca Veta La Palma (Isla Mayor, Spain).

### 2.3. *In vivo* dose-response of rFsh on female steroid production

To evaluate the biological potency of rFsh produced in CHO cells in inducing 17β-estradiol (E2) production and to determine the minimum effective dose and optimal dosing schedule, a single intramuscular injection of different rFsh doses (0, 3, 6, 9, 12 and 15 μg kg−1) was administered to immature grey mullet females (five fish per dose group) (mean body weight 0.9 ± 0.3 kg). Blood samples (0.40 mL) were collected before injection (day 0) and at different days (1, 3, 6, 9, 13, 17, 21 days) after injection. Control females were injected with CHO conditioned culture medium (1 mL fish-1).

### 2.4. Experiment 1. Long-term rFsh therapy

In Experiment 1, twenty-six adult flathead grey mullet were used in the trial. Nine females and three males (mean ± SD body weight 1 ± 0.3 and 0.9 ± 0.1 kg; mean standard body length 41.4 ± 4.1 and 40.8 ± 2.4 cm, respectively) to the gonadotropic treatment group and 11 females and three males (mean ± SD body weight 1 ± 0.2 and 0.9 ± 0.1 kg; mean standard body length 42 ± 4.1 and 41.3 ± 1.5 cm, respectively) to the control group. The total biomass was 24.7 kg. Only three males were selected for each group, as only six males were available. The fisheries capture to form the broodstock was biased towards females as has been observed in other studies (Rao and Babu, 2016). The fish in the treatment group were administered rFsh followed by either hCG alone (El-Gharabawy and Assem, 2006; Yousif et al., 2010) or GnRH combined with DA antagonist (Aizen et al., 2005).

#### 2.4.1 Stage 1. Induction of gametogenesis in males and females

Individuals belonging to the gonadotropic treatment group (both males and females) received weekly intramuscular injections of specific grey mullet rFsh at a dose of 15 μg kg−1 for 11 weeks (Fig 1). The rFsh dose applied was chosen according to the dose with highest potency on E2 induction in the in vivo dose-response study. The dose and the time frame of administration were also selected based on the results obtained in a previous study on Senegalese sole (Solea senegalensis) using recombinant Gths produced in CHO cells. Chauvigné et al. (2017) described that a dose of 12 - 17 μg kg−1 rFsh was effective in stimulating spermatogenesis, while the hormone was detectable in the bloodstream for approximately seven days. The control fish were injected in the same manner as rFsh treated fish, but with CHO conditioned culture medium (1 mL fish-1). Fish were sampled before the first injection and on different weeks before receiving the corresponding weekly injection. At fortnightly intervals, blood samples (0.40 mL) from the caudal vein and oocytes through cannulation were obtained. The diameter of the largest oocytes (n = 20) per female were measured in situ and samples were fixed for histology. In parallel, males received a gentle abdominal pressure to check the presence of milt.

**Figure 1.**
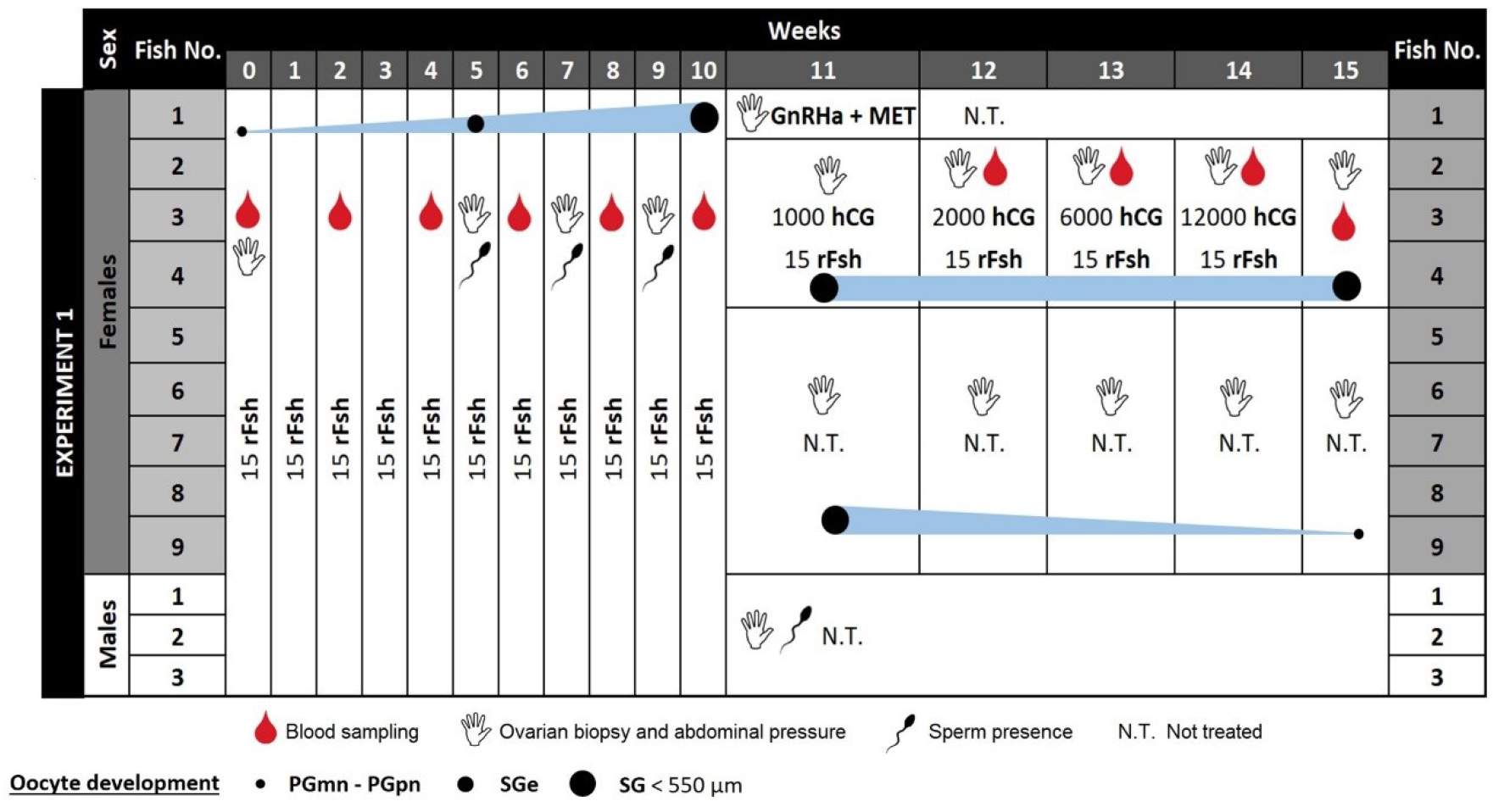
Schematic representation of the induction protocol administered to flathead grey mullet (M. cephalus) in Experiment 1. Columns represent weeks of each experiment and rows represent the different fish. Adult grey mullet females (n = 9) and males (n = 3), received weekly doses of intramuscular injections of rFsh. From 11 weeks onwards, the females with the most advanced stages of vitellogenesis received different weekly treatments. Female 1 received a GnRHa + MET protocol consisted of a priming (GnRHa 10 μg kg−1; MET 15 mg kg−1) and a resolving (GnRHa 20 μg kg−1; MET 15 mg kg−1) injection administered 22.5 h apart (Aizen et al., 2005), whilst females 2 - 4 were administered increasing doses of hCG in addition to rFsh. Control individuals (n = 11 females, n = 3 males) received weekly injections of CHO conditioned culture medium (1 mL fish-1) during 11 weeks. Doses of rFsh and rLh are expressed in μg kg−1 and doses of hCG in IU kg−1. A hand symbol represents when ovarian biopsies or abdominal massage for sperm were made, red drops represent blood sampling, a spermatozoa represents when males had flowing sperm and circles represent the different stages of oocyte development encountered; small black is primary growth, medium black is early secondary growth (vitellogenesis), large black is advanced secondary growth < 550 μm oocyte diameter.

#### 2.4.2 Stage 2: Completion of oocyte growth and maturation induction in females

This second stage of the experiment investigated the effects of different hormones used as a source of Lh or to induce endogenous Lh release to complete oocyte growth and induce maturation in adult females that were treated with rFsh to induce vitellogenesis. Five females were not used in the second stage and rFsh administration was stopped, although oocyte changes were assessed until the end of the experiment. Stage 2 focused on the four fish with the most advanced stages of vitellogenesis. One female with a mean maximum oocyte diameter of 539 ± 5 μm was treated with the GnRHa des-Gly10, [D-Ala6]-gonadotropin releasing hormone (product code L4513, Sigma, Spain) in combination with Metoclopramide (MET) (product code M0763, Sigma, Spain), a dopamine antagonist, according to the Aizen et al. (2005) protocol, which consisted of a priming (GnRHa 10 μg kg−1; MET 15 mg kg−1) and a resolving (GnRHa 20 μg kg−1; MET 15 mg kg−1) injection administered 22.5 h apart. Three females that reached a mean maximum diameter of 450 ± 10, 450 ± 9 and 470 ± 8 μm after rFsh treatment, received weekly consecutive injections of hCG (Veterin Corion, DIVASA-FARMAVIC S.A, Barcelona) at increasing doses (1000, 2000, 6000, 12000 IU kg−1) in combination with the rFsh treatment (15 μg kg−1) (Fig 1). Dosage of hCG were in the range of previous studies on grey mullet maturation (El-Gharabawy and Assem, 2006; Yousif et al., 2010) and other fish species (Mañanós et al., 2009). Weekly samples of oocytes and blood (0.40 mL) were obtained.

### 2.5. Experiment 2. Combined rFsh and rLh therapy

A total of twenty-four adult flathead grey mullet were used in Experiment 2. Females had a body weight of 0.9 ± 0.1 kg (mean ± SD) and standard length of 38.5 ± 3.1 cm, and males 0.6 ± 0.1 kg and 33.3 ± 1.2 cm formed the treatment group, while females with a body weight of 0.8 ± 0.1 kg and standard length of 39.5 ± 1.3 cm and males with 0.8 ± 0.1 kg and 38.6 ± 2.7 cm were used as controls. The total biomass was 20 kg. In this trial, 2/3 of the females were immature with previtellogenic ovaries, while 1/3 were at the beginning of secondary growth (with cortical alveoli stage oocytes). All females were randomly distributed between treated and control groups. The aim of the administration pattern was to simulate natural increases and decreases of gonadotropins in the bloodstream of individuals according to their suggested regulatory role in gamete development (Levavi-Sivan et al., 2010). Initial administration of rFsh followed by a gradual increase of rLh as gametogenesis progresses and subsequent decline of rFsh.

#### 2.5.1. Females

Initially, all nine females received increasing doses of rFsh, 6 μg kg−1 (week 0) and 9 μg-1 kg (week 1) before the dose was fixed at 12 μg kg−1 rFsh per week (Fig 2). A maximum 12 μg kg−1 dose was selected for long-term treatment based on Experiment 1 and the in vivo dose-response study. From the 4th week onwards, females (n = 8) were also administered a weekly injection of rLh at increasing doses (2.5, 4, 6 μg kg−1). When vitellogenesis arrived to advanced stages (week 9), weekly rFsh dose was decreased to 4 μg kg−1 while rLh dose was increased (9 and 12 μg kg−1). At this point (week 8 and onwards), treatments were adjusted accordingly to oocyte diameter of each individual fish. When females presented oocytes ≥ 550 μm, oocytes were considered to have completed vitellogenic growth, therefore, no more rFsh was administered and consecutive doses starting with 9 and maintaining 12 μg kg−1 rLh were administered every 3 days. The aim of this increased frequency of administration was to maintain high levels of rLh in the bloodstream, based on the half-life (shorter than rFsh) described for rLh produced in CHO cells and administered to Senegalese sole (Chauvigné et al., 2017). When the most developed oocytes reached a diameter ≥ 600 μm or did not show further growth, females were administered higher doses of rLh (15 or 30 μg kg−1) combined with a 40 mg kg−1 intramuscular injection of progesterone (Prolutex, IBSA Group, Italy) administered 24 h after the rLh injection to induce oocyte maturation, ovulation and spawning (see schematic representation of the experimental setup in Fig 2). Three females received 15 μg kg−1 of rLh and five females received 30 μg kg−1.

**Figure 2.**
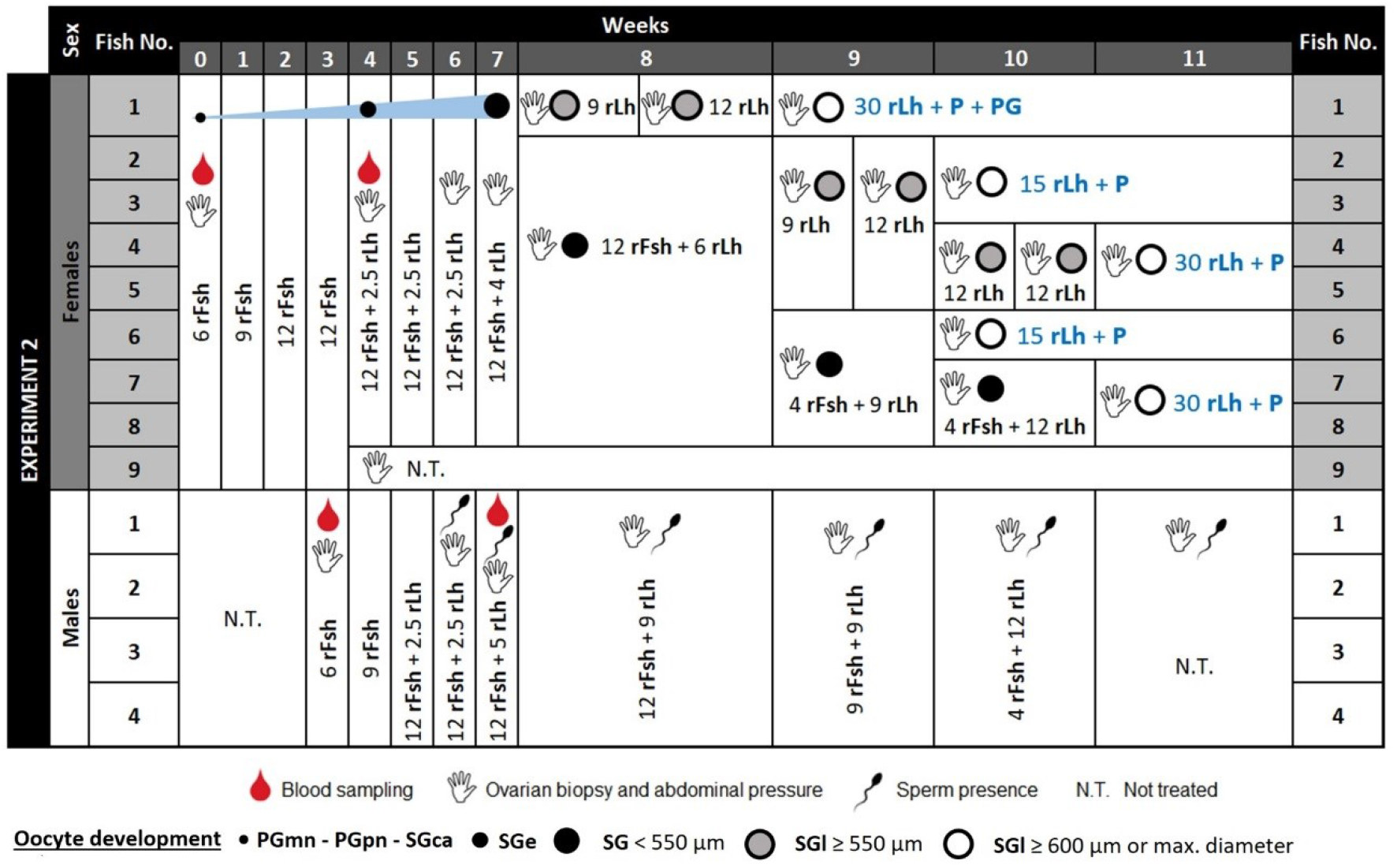
Schematic representation of the induction protocol administered to flathead grey mullet (M. cephalus) in Experiment 2. Columns represent weeks of each experiment and rows represent the different fish. Females (n = 9) received increasing doses of rFsh, and from the 4th week combined with increasing doses of rLh, followed by a decrease in rFsh. When females presented ≥ 550 μm oocytes rLh was administered every three days (weeks 9 and 10 for females 2 - 5). When the most developed oocytes reached a diameter of ≥ 600 μm, females were administered higher doses of rLh 15 μg kg−1 (females 2, 3 and 6) or 30 μg kg−1 (females 1, 4, 5, 7 and 8) combined with a 40 mg kg−1 injection of P administered 24 h after the rLh injection to induce oocyte maturation, ovulation and spawning. Female 1 was also administered 18.75 μg kg−1 of prostaglandin F2α (PG) 39 hours after the rLh injection. Males (n = 4) initiated rFsh treatment on week 3 and were administered a similar, but shortened program of increasing rFsh dose followed with a combined increasing rLh before decreasing rFsh. The individuals in control groups (n = 7 females, n = 4 males) underwent the same number of intramuscular injections as treated individuals, but with CHO culture medium (1 mL fish-1). Doses of rFsh and rLh are expressed in μg kg−1 and doses of P in mg kg−1. A hand symbol represents when ovarian biopsies or abdominal massage for sperm were made, red drops represent blood sampling, a spermatozoa represents when males had flowing sperm and circles represent the different stages of oocyte development encountered; small black is primary growth to cortical alveoli stage, medium black is early secondary growth (vitellogenesis), large black is advanced secondary growth < 550 μm oocyte diameter, large grey is late secondary growth ≥ 550 μm diameter and large white indicate full-grown secondary-growth oocytes of ≥ 600 μm diameter or maximum diameter achieved.

After the application of rLh, females were placed in a 10 m3 tank with spermiating males. Surface out-flow egg collectors were placed to receive eggs from the tanks and were checked for eggs regularly. The fish were also observed frequently (from outside of the tank), for swelling of the abdomen (hydration) in females and the initiation of courtship behaviour. These frequent checks were made as there is no established latency time of spawning for rGth treatments. Latency time reported for grey mullet after resolving doses from other hormone treatments varies from 17 to 48 hours at 22 - 25 °C (El-Gharabawy and Assem, 2006; Yousefian et al., 2009). One female (female 1, 30 μg kg−1 rLh + 40 mg kg−1 P in Fig 2) that had oocytes ≥ 600 μm earlier (week 9) than the other females, developed a large swollen belly without ovulation and was administered 18.75 μg kg−1 of prostaglandin F2α (VETEGLAN, Laboratorios Calier, S.A., Spain) 39 hours after the rLh injection. The other seven females (females 2 - 8 in Fig 2) did not receive prostaglandins and were checked and/or stripped as there was no natural spawning. Four females ovulated and were stripped, one female (female 1) at 40 h and three (females 4, 5 and 7) at 48 ± 0.5 h after the rLh injection. Total number of eggs (fecundity) was estimated by counting the number of eggs in triplicate in a subsample of 500 μl.

The seven females in the control group underwent the same number of intramuscular injections as treated females but with CHO culture medium (1 mL fish-1). Females were sampled for oocyte tissue (weeks 0, 4, 6, 7, 8, 9, 10, 11, immediately before hormone administration) and blood (week 0 – before treatment, week 4 – after 4 weeks of rFsh treatment).

#### 2.5.2. Males

The treatment of males in the rGth group (n = 4) initiated three weeks after the females in order to synchronise development of both sexes and have sperm and eggs available at the same time for fertilisation. The same rFsh doses were applied as for females and the dose range of rLh was fixed accordingly to other studies in male spermatogenesis and spermiogenesis (Chauvigné et al., 2018; Peñaranda et al., 2018) (Fig 2).

The four males in the control group were treated as previously reported for control groups. Males were sampled for sperm (weeks 3, 6, 7, 8, 9, 10 and 11 of the experiment) and blood (week 3 – before treatment, week 7 – after 4 weeks of hormone treatment).

#### 2.5.3. *In vitro* fertilisation

For the in vitro fertilisation, sperm was obtained from three males, diluted 1:4 in the extender solution Marine Freeze® (IMV Technologies, France) that showed the best results for sperm conservation in a marine species (González-López et al., 2020) and stored at 4°C to fertilise the eggs. The eggs from each female (n = 3) were stripped and total volume registered. Aliquots of 0.5 mL of eggs (~1200 eggs) from each female were each fertilised in triplicate with a pool of 60 μL of diluted sperm (20 μL from each of the three males, ~190,000 spermatozoa egg−1) (3 females × 3 triplicates = 9 fertilisations). The diluted sperm was pipetted directly onto the 0.5 mL of eggs in a 100 mL beaker and immediately activated by mixing the eggs and sperm with 5 mL of clean tank water. After 5 minutes, the beaker was filled to 100 mL with clean tank water and placed in a temperature-controlled incubator (24°C) to incubate the eggs. Twenty-two hours after fertilisation, all eggs were checked for embryo development and the percentage of eggs fertilised was calculated as the number of eggs with live embryos/number of eggs used for the in vitro fertilisation. Eggs with embryonic development were transferred individually into individual wells filled with sterile seawater in a 96 well plate and incubated (24°C). To evaluate the quality of the eggs with embryo, the hatching success was calculated as the number of hatched larvae/live 22-hours embryos. Larvae was checked every 24 h until all hatched larvae had died and larval survival rate was calculated as the number of live larvae / larvae that hatched. A subsample of ~1/3 fertilised eggs and larvae were used for taking measurements and afterwards returned to the incubation.

### 2.6. Plasma steroid analysis

Blood samples were centrifuged at 3,000 rpm at 4 °C for 15 min and the plasma stored at −80 °C until steroid analysis. Plasma levels of E2 and 11-ketotestosterone (11-KT) were measured for females and males, respectively, and were analysed using a commercially available enzyme immunoassay (EIA) kits (Cayman Chemical Company, USA). Steroids were extracted with methanol, which was evaporated and extracts were re-suspended 1:10 in the EIA buffer.

### 2.7. Histological observations and classification of developing ovaries

Ovarian biopsy samples were preserved in Bouin’s fluid, dehydrated through an ethanol series and embedded in paraffin. Histological sections (3 μm) were stained with hematoxylin and eosin (Casa Álvarez, Spain). To examine ovarian development, oocytes sections were observed under a light microscope (Leica DMLB, Houston, USA). Quantification of the percentage of oocytes in different stages in the ovaries among weeks was made by the identification of 50 - 100 random oocytes per female each week. Oocyte developmental stage was based on the identification of structures, morphological changes and increasing oocyte diameter. Oocytes were classified as: multiple nucleoli step of primary growth (PGmn) characterised by small oocytes with multiple nucleoli that were not situated in the periphery of the germinal vesicle, perinucleolar step of primary growth (PGpn), with the nucleoli located around the internal germinal vesicle membrane, cortical alveoli step (SGca), determined by the presence of small oil droplets and granular vesicles “cortical alveoli” in the peripheral ooplasm, early secondary growth (SGe), with the appearance of yolk globules and with this the initiation of vitellogenesis, secondary growth (SG) corresponding to mid-to late-vitellogenesis when oocytes reached ≥ 400 μm (Greeley et al., 1987), oocyte maturation stage (OM), with the identification of coalesced oil droplets and the displacement of the germinal vesicle to the ooplasm periphery and some hydration and coalescence of yolk globules, and ovulation stage (OV), when one large yolk globule is observed (Lubzens et al., 2010). Atresia was identified by the hypertrophy of granulosa cells, the loss of the individuality of yolk globules and the dissolution of their content (Valdebenito et al., 2011).

### 2.8. Sperm collection and evaluation

Sperm samples were collected in a 1 mL syringe avoiding the contamination by faeces, urine and / or sea water. Approximately 1 μL of sperm was placed on a microscope slide beside 0.2 mL of sea water, mixed to activate the spermatozoids and immediately (first 10 seconds) observed through a microscope at 100x magnification (Zeuss Microscopes). The assessment of the milt quality was estimated by the percentage of motile spermatozoa and by the total duration of the movement from sperm activation until all forward movement of spermatozoa stopped. The observations were made in triplicate and the percentage of motile spermatozoa was classified into different motility scores: 0 for no motile sperm, 1 for > 0 – 25 % of sperm with progressive movement, 2 for > 25 % - 50 % of sperm with progressive movement, 3 for > 50 – 75 % and 4 for > 75 % of sperm with progressive movement (Mañanós et al., 2009). For those samples in Exp. 2 with a motility score of 4 and manageable sperm volumes (≥ 100 μL) (n = 10), sperm quality was also evaluated using a CASA system (Wilson-Leedy and Ingermann, 2007). For this, 0.5 μL of diluted sperm (1/4 in Marine Freeze®) were dropped on the centre of a slide and activated using 20 μL of sea water. A 1 μL sample containing the activated spermatozoa was pipetted into an ISAS counting chamber (Integrated Sperm Analysis System, Spain). The tracks of the activated spermatozoa were recorded through a bright field equipped video microscope at 200x magnification (Olympus BH Microscope and DMK 22BUC03 Camera with 744×480 “0.4 MP” resolution at 60 FPS, The Imaging Source Europe GmbH, Bremen, Germany). The video sections from 15 to 17 s after activation were transformed to image sequences using VIRTUALDUB 1.9.11 (virtualdub.org) free software. The spermatozoa in each field were selected by adjusting the grayscale threshold through Image J software (https://imagej.nih.gov/ij/). The following sperm quality parameters were determined: (1) sperm motility (%), (2) sperm velocity (μm s−1): the curvilinear velocity (VCL), straight-line velocity (VSL) and average path velocity (VAP), (3) sperm movement trajectory: path linearity of actual sperm track, LIN = VSL/VCL × 100), path wobble (deviation from average path, WOB = VAP/VCL × 100), and path straightness (linearity of the average path, STR = VSL/VAP × 100). All parameters were evaluated in triplicate for each sperm sample.

Sperm concentration was also recorded for each sperm sample used. In this case, sperm was diluted 1/1000 and 10 μL were pipetted into a THOMA cell counting chamber where it was allowed to settle for 10 min, and then, was observed under the microscope at 100x magnification. The estimated densities are expressed as the number of spermatozoa per mL of sperm (spz mL−1). Quantification of spermatozoa was conducted using ImageJ software.

### 2.9. Statistical analysis

Shapiro-Wilk and Levene tests were used to check the normality of data distribution and variance homogeneity, respectively. Oocyte diameter data (Stage 1 from Exp. 1 and Exp. 2), E2 levels (Stage 1 from Exp. 1 and 2) and 11-KT levels (Exp. 1) were normalised with the ln log transformation. For oocyte diameter, E2 levels and 11-KT levels (Stage 1 from Exp. 1 and Exp. 2) a two-way repeated-measures (RM) ANOVA followed by Dunnett’s test was used to compare to the control, which was the control group and week 0 of treatment. A t-student was used to compare oocyte diameter before and after the Stage 2 treatments in Experiment 1. Differences in weekly E2 levels in Stage 2 (Exp. 1) treatments were examined by one-way RM ANOVA. Statistical differences in the dose-response test and in sperm characteristics (density, duration) among weeks were examined by a one-way repeated-measures analysis of variance (ANOVA) followed by the Holm-Sidak test for pairwise comparisons. Analyses were performed using SigmaPlot version 12.0 (Systat Software Inc., Richmond, CA, USA). Significance was set at P < 0.05. Data is presented as mean ± standard error (SEM) unless indicated otherwise.

## 3. Results

### 3.1. *In vivo* dose-response of rFsh on female steroid production

There were no increases from the E2 basal values after the application of doses of 3, 6 and 9 μg kg−1 of rFsh (Fig 3). A great individual variation in magnitude of response was observed when a dose of 9 μg kg−1 was administered. The administration of 12 μg kg−1 of rFsh produced significant increases in E2 levels on 3 to 6 days after the injection. The highest average levels of E2 were obtained 3 days after the injection of 15 μg kg−1. Therefore, the doses of 12 to 15 μg kg−1 of rFsh were the most effective to stimulate E2 production and were considered the most appropriate for the induction experiments.

**Figure 3.**
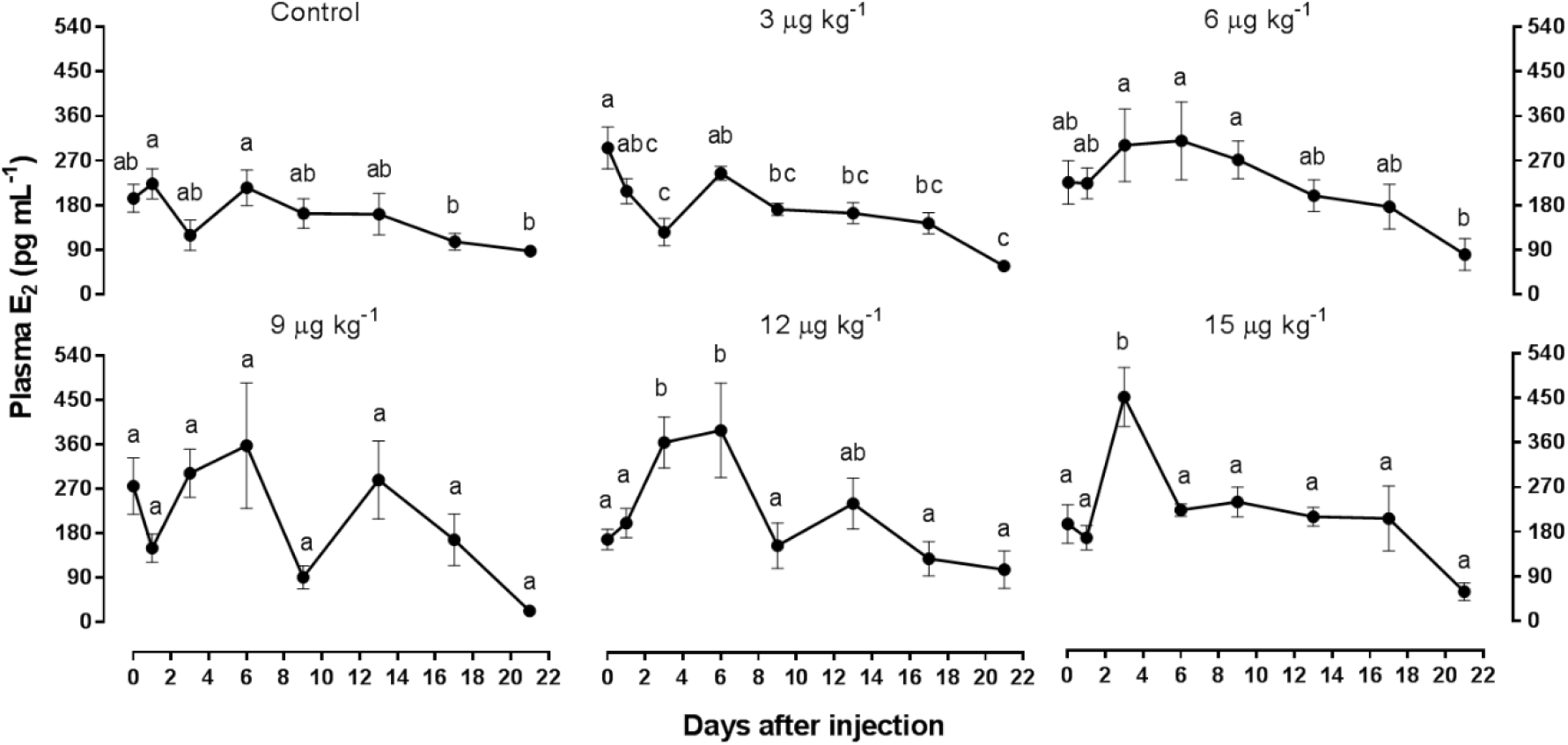
Mean (± SEM) plasma E2 levels of female flathead grey mullet (M. cephalus) before (day 0) and after (day 1, 3, 6, 9, 13, 15 and 21 days) the rFsh injection. Females (n = 5/group) received a single injection of rFsh at doses 3, 6, 9, 12 or 15 μg kg−1 and an injection of 1 mL fish-1 CHO conditioned culture medium for control. Different letters indicate significant differences (P < 0.05) over time within each dose.

### 3.2. Experiment 1: Effect of long-term rFsh therapy in female development

#### 3.2.1. Stage 1: Gametogenesis induction in females

Weekly injections of 15 μg kg−1 rFsh during eleven weeks to immature females generated a significant increase (2 - 10 weeks) in the plasma levels of E2 compared to the control group (P < 0.001) (Fig 4). Among the untreated females (control), plasma E2 levels remained unchanged at basal levels during the experimental period (0 - 10 weeks). In situ and histological observation of oocytes obtained by cannulation indicated that rFsh administration induced a significant increase of oocytes diameter (P < 0.001) (Fig 5A) and vitellogenic growth (Fig 6) compared to the control group.

**Figure 4.**
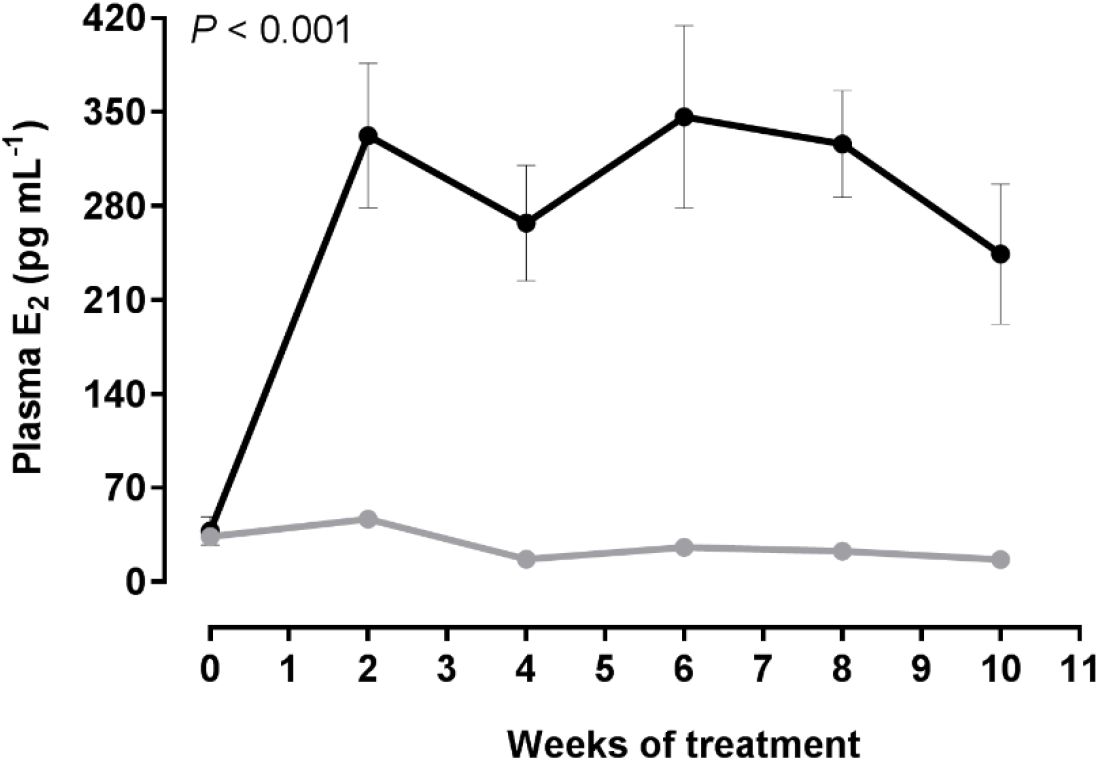
Mean (± SEM) plasma E2 levels of rFsh-treated and control flathead grey mullet (M. cephalus) females (n = 9-11) in Experiment 1. Treated females received weekly injections of rFsh (15 μg kg−1) and control females of CHO conditioned culture medium (1 mL fish-1). There were significant differences among treatments (two-way repeated measures ANOVA, P < 0.001)

**Figure 5.**
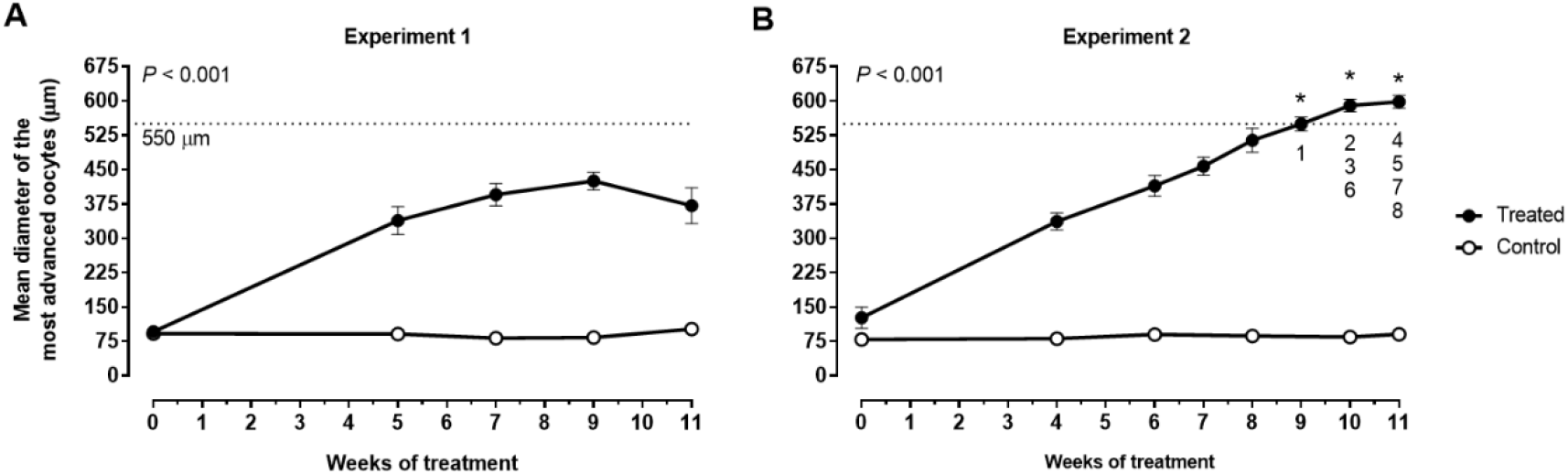
Mean (± SEM) oocyte diameter of the most developed oocytes in wet mounts from rFsh treated and control flathead grey mullet (M. cephalus) females. (A) Experiment 1, females treated (n = 9) with a weekly 15 μg kg−1 rFsh administration or CHO conditioned culture medium (control, n = 11) during 11 weeks. (B) Experiment 2, females treated (n = 9) with initial increasing doses of rFsh followed by increases in rLh and subsequent rFsh decrease or CHO conditioned culture medium (control, n = 7). Values used for females checked twice in the same week were the mean of both revisions. Asterisks show the moment when numbered females (see Fig 2) were selected for maturation and ovulation induction. There were significant differences between treated and control groups (two-way repeated measures ANOVA, P < 0.001). Dotted line indicates oocyte size recommended for the hormonal induction of oocyte maturation.

**Figure 6.**
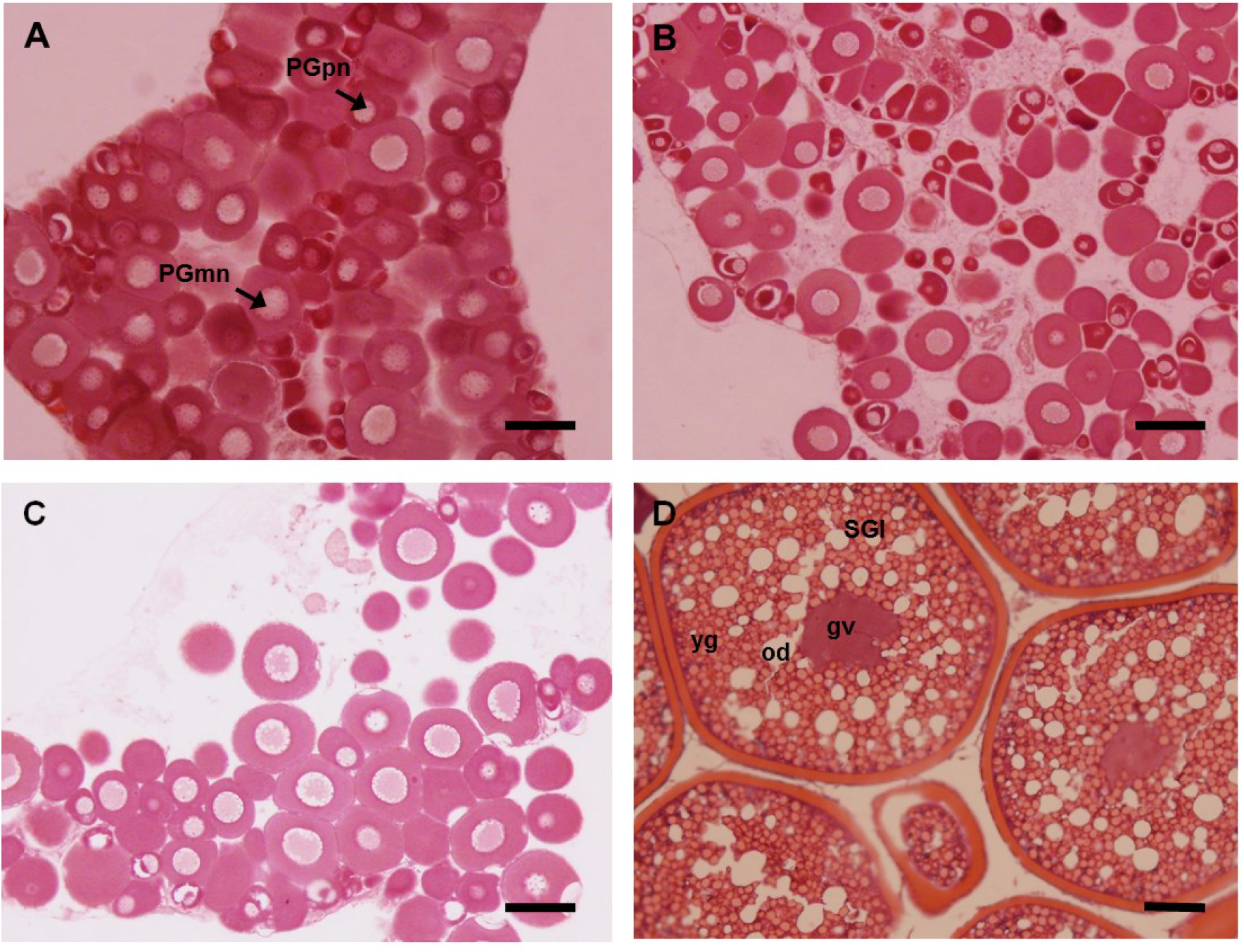
Effects of long-term treatment of rFsh on ovarian development in previtellogenic flathead grey mullet (*M. cephalus*) *in vivo*. Histological sections stained with hematoxylin and eosin show oocytes samples from (A) initial control fish, (B) rFsh-treated fish before treatment, (C) control fish after 7 weeks and (D) rFsh-treated fish after 7 weeks of treatment (weekly 15 μg kg−1 rFsh). gv, germinal vesicle; od, oil droplets; PGpn, perinucleolar primary growth oocyte; PGmn, multiple nucleoli primary growth oocyte; SGl, late secondary growth oocyte; yg, yolk globules. Scale bar: 100 μm.

At the beginning of the treatment all females presented oocytes at the PGpn (mean maximum diameter = 97 ± 4 μm) (Fig 6) with the exception of one female assigned to the rFsh-treated group that presented oocytes at PGmn. After 5 weeks of treatment, all rFsh-treated females (89%) except one had vitellogenic oocytes (Fig 7A). In addition, some traces of atresia appeared in some females. In the two subsequent revisions (weeks 7 and 9), SG oocytes were the most abundant with a maximum size of 425 ± 19 μm in diameter (Fig 5A). After 9 weeks of treatment, the proportion of atresia observed in the vitellogenic ovaries increased from 3 to 24 % (Fig 7A). The female that at the start of the experiment before any treatment had oocytes at PGmn was delayed compared to other females and only developed to SGe after 11 weeks of treatment. Therefore, of the nine treated females all (100%) developed from pre-vitellogenic oocyte stages to vitellogenesis and eight (89%) developed to late vitellogenic stages of oocyte development. In comparison, the oocytes of all (100%) untreated females remained at primary growth (perinucleolar stage) during the entire experiment (Figs 5A, 6A). When rFsh administration for five females was ceased from week 11 onwards, the morphology of the ovary returned to primary growth oocytes after five weeks.

**Figure 7.**
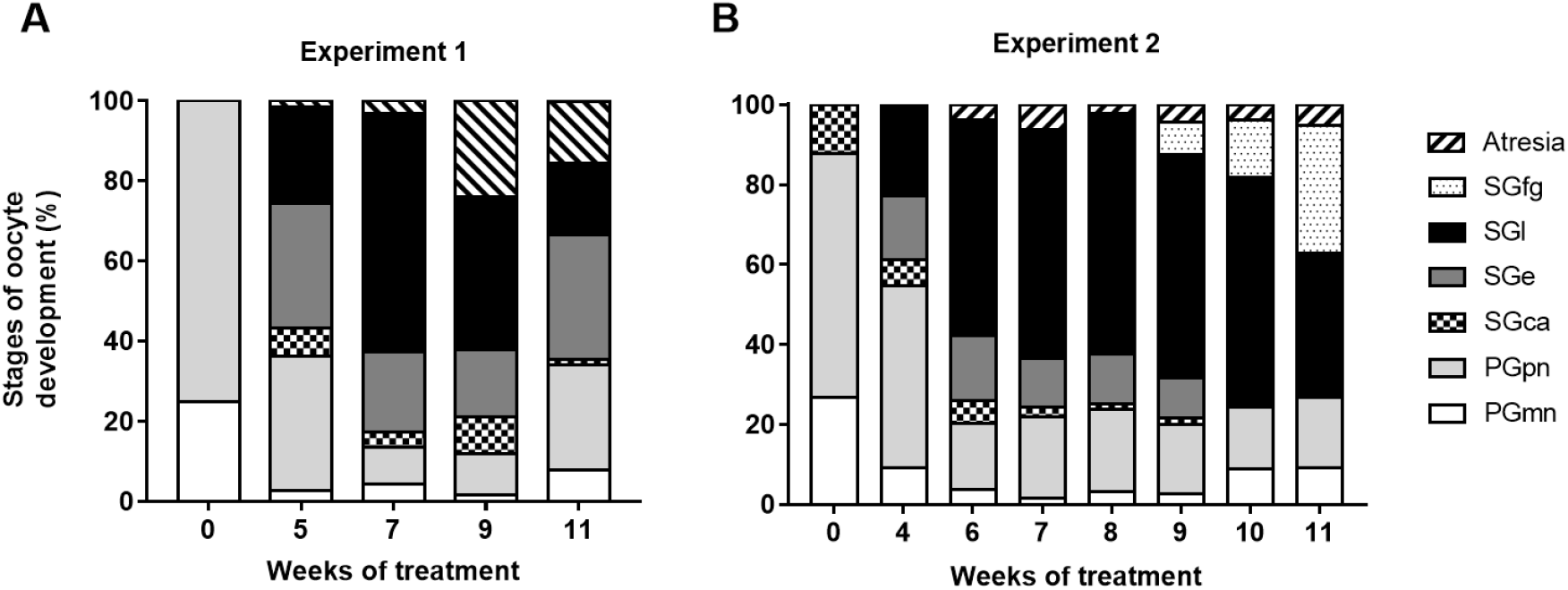
Temporal weekly evolution of percentage frequency of oocyte developmental stages observed in rFsh treated flathead grey mullet females (M. cephalus). (A) Experiment 1 with weekly 15 μg kg−1 rFsh administration to the treated group (n = 9) during 11 weeks. (B) Experiment 2 with the administration of initial increasing doses of rFsh followed by increases in rLh and subsequent rFsh decrease (n = 9). PGmn, multiple nucleoli step of primary growth; PGpn, perinucleolar primary growth oocyte; SGca, cortical alveoli step; SGe, early secondary growth; SGl, late secondary growth oocyte; SGfg, full-grown secondary-growth oocytes.

#### 3.2.2. Stage 2: Completion of oocyte growth and maturation

Histological examination of the oocytes after each treatment (GnRHa+MET or hCG) did not show variations in oocyte morphology although a significant increase in oocyte diameter was observed in the female injected with GnRHa+MET protocol (Table 1). The injections of hCG at doses of 1000, 2000, 6000, 12000 IU kg−1 combined with 15 μg kg-1 rFsh did not completed oocyte growth and oocyte maturation. High E2 levels were maintained during the period of weekly hCG injection (week 12: 186.5 ± 20.6, week 13: 258.3 ± 35.1, week 14: 241.1 ± 42.1 and week 15: 184.5 ± 30.8 pg mL−1) that were not significantly different from E2 levels (391.4 ± 56.5 pg mL−1) during weeks 4 - 10 (Stage 1) in the same group.

**Table 1.** Effects of treatments applied to flathead grey mullet (*M. cephalus*) to induce completion of oocyte growth and oocyte maturation in Stage 2 from Experiment 1. Differences (t-student, P < 0.05) between maximum oocyte diameter (mean ± SEM) reached with rFsh treatment at Stage 1 and final oocyte diameter after corresponding treatments are indicated by different letters for each female.

| Fish No. | Max. oocyte diameter reached with rFsh at Stage 1 ($\mu\text{m}$) | Priming GnRHa ($\mu\text{g kg}^{-1}$); MET ($\text{mg kg}^{-1}$) | Resolving GnRHa ($\mu\text{g kg}^{-1}$); MET ($\text{mg kg}^{-1}$) | Weekly rFsh ($15 \mu\text{g kg}^{-1}$); hCG ($\text{IU kg}^{-1}$) | Final max. oocyte diameter at Stage 2 ($\mu\text{m}$) |
| --- | --- | --- | --- | --- | --- |
| 1 | $539 \pm 5^a$ | 10; 15 | 20; 15 | - | $569 \pm 10^b$ |
| 2 | $450 \pm 10^a$ | - | - | 1000, 2000, 6000, 12000 | $437 \pm 6^a$ |
| 3 | $450 \pm 9^a$ | - | - | 1000, 2000, 6000, 12000 | $422 \pm 8^b$ |
| 4 | $470 \pm 8^a$ | - | - | 1000, 2000, 6000, 12000 | $490 \pm 8^a$ |

### 3.3. Experiment 2: Effect of combined rFsh and rLh therapy in female development

Histology showed that at the beginning of the treatment, those females from semi-extensive culture and with less time in intensive captive conditions (3 months) had oocytes in cortical alveoli stage (2 were in the control group and 3 in the gonadotropin treated group, Table 2), while the wild individuals with 12+ months in captivity had primary growth perinucleolar stage as the most advanced stage of gonadal development (5 in control group and 6 in treated group, Table 2).

**Table 2.** Individual flathead grey mullet (*M. cephalus*) oocyte diameter before inducing oocyte maturation (mean ± SEM), hormone treatment and egg fecundity data in Experiment 2. PGpn, perinucleolar primary growth oocyte; SGca, cortical alveoli step.

| Fish No. | Oocyte stage at the start of the experiment | Max. oocyte diameter before OM induction ($\mu\text{m}$) | rLh ($\mu\text{g kg}^{-1}$) (t = 0) | Progesterone ($\text{mg kg}^{-1}$) (t = 24) | Prostaglandins ($\mu\text{g kg}^{-1}$) (t = 39) | Total eggs | Fecundity (eggs $\text{kg}^{-1}$ bw) | Fertilisation rate (%) |
| --- | --- | --- | --- | --- | --- | --- | --- | --- |
| 1 | SGca | $625 \pm 8$ | 30 | 40 | 18.75 | 801,913 | 832,723 | Not used |
| 2 | PGpn | $619 \pm 7$ | 15 | 40 | - | - | - | - |
| 3 | SGca | $627 \pm 8$ | 15 | 40 | - | - | - | - |
| 4 | PGpn | $603 \pm 10$ | 30 | 40 | - | 974,928 | 574,500 | 0.1 |
| 5 | SGca | $608 \pm 8$ | 30 | 40 | - | 754,774 | 676,320 | 0.31 |
| 6 | PGpn | $610 \pm 6$ | 15 | 40 | - | - | - | - |
| 7 | PGpn | $578 \pm 7$ | 30 | 40 | - | 891,600 | 888,047 | 0.81 |
| 8 | PGpn | $605 \pm 4$ | 30 | 40 | - | - | - | - |

As in Experiment 1, the administration of rFsh significantly (P < 0.001) increased the production of E2 (week 0: 123.9 ± 27.4; week 4: 458.7 ± 113 pg mL−1) compared to the control group (week 0: 95.6 ± 21.5; week 4: 81.1 ± 18.7 pg mL−1). This increase in E2 levels in Exp 2, was achieved despite of using a lower and increasing dose during the first weeks (Fig 2). After the first 4 weeks of treatment, all but one female (89 %) had vitellogenic oocytes. The treatment of the delayed non-vitellogenic female (female 9 in Fig 2) was stopped, even though the diameter of the most developed oocytes had increased significantly from week 0 (89 ± 2 μm) to week 4 (167 ± 3 μm). Oocyte growth of all other females followed the same pattern as observed in Exp 1 during the first 7 weeks of treatment (Fig 7). However, during the following weeks, with the administration of rLh, the proportion of atresia was reduced (week 9 = 4%) in comparison with Exp 1 (24%) in which just rFsh was administered. The inclusion of rLh in Exp 2 also increased the mean diameter of the most advanced oocytes compared to Exp 1 (Fig 5A vs 5B). The response to treatment was different between females, which reached a ≥ 550 μm oocyte diameter at different time points between week 8 and 11. Full-grown oocytes were obtained in all eight (89%) females and prior to maturation induction had a mean diameter of 609 ± 5 μm. As in Exp 1, all (100%) control females showed no oocyte growth or development.

The three females that received 15 μg kg−1 rLh followed by 40 mg kg−1 of progesterone did not respond to the treatment and no significant increase in oocyte diameter or developmental stages was observed. Only those females (n = 5) that received 30 μg kg−1 of rLh followed by 40 mg kg−1 of progesterone, presented oocyte maturation (OM), hydration and ovulation. Five females showed the initiation of OM indicated by oil globule coalescence after 24 h from rLh injection. From these females, female 1 had not ovulated 39 hours after rLh administration when an injection of prostaglandin F2α was administered. The prostaglandin appeared to induce ovulation and, one hour after administration, poor quality eggs were stripped that were not used for fertilisation. Posterior histological analysis showed that the eggs were not fully hydrated (Fig 8A and Table 2). Three females, females 4, 5 and 7, which were checked at 48 ± 0.5 hours from rLh injection, ovulated (Fig 8B) and after stripping, eggs were used for in vitro fertilisation (Table 2). The mean relative fecundity was 742,900 ± 71,840 eggs kg−1 bw. Female 8 did not ovulate and at 48 ± 0.5 hours after rLh administration only presented oocytes in OM. Therefore, of the nine females, eight (89%) terminated vitellogenesis to stage immediately prior to OM, five (56%) were induced with 30 μg kg−1 of rLh + P and 100% of these five advanced to OM, four (80%) ovulated and three of them (60%) had a low percentage of viable eggs according to fertilisation rates.

**Figure 8.**
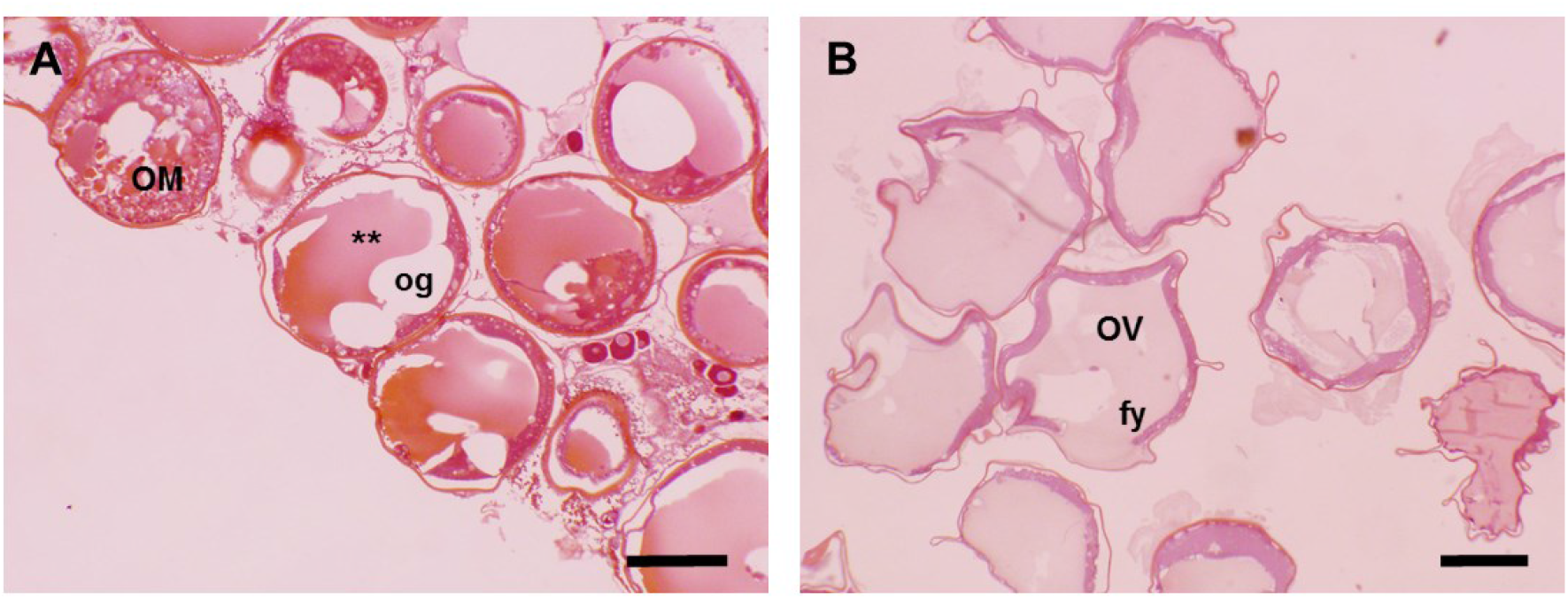
Oocyte maturation and hydration stages for treated flathead grey mullet females (Mugil cephalus) in Experiment 2. (A) Ovulated eggs from female 1 at 40 hours after 30 μg kg−1 of rLh injection (16 hours from 40 mg kg−1 progesterone) and 1 hour from 18.75 μg kg−1 prostaglandin F2α injection. Oocytes in maturation: yolk globules coalesce and fuse to form a one large globule (**). Central oil globule displaces the germinal vesicle into an eccentric position. (B) Ovulated eggs from three females (females 4, 5 and 7) at approx. 48 hours after 30 μg kg−1 of rLh injection (24 hours from 40 mg kg−1 progesterone). Oocytes have undergone hydration after completion of germinal vesicle breakdown with homogenous fluid yolk. fy, fluid yolk; og, oil globules; OM, oocyte maturation; OV, hydrated oocytes at ovulation stage. Scale bar: 500 μm.

### 3.4. Male development

Control males did not produce milt neither in Experiment 1 (n = 3) nor in Experiment 2 (n = 4). In Experiment 1, in the first revision after five weeks of rFsh treatment, two of three (66.6%) males produced sperm that coincided with an increase in 11-KT levels (P = 0.043, α = 0.05, statistical power = 0.66) (Fig 9). The production of sperm was prolonged for 6 weeks, but sperm was highly viscous and sperm volumes were low (29.3 ± 7.1 μL), which made it difficult to manipulate. The mean sperm concentration was 4.6 ± 1.5 1010 spermatozoa mL−1, the motility grade recorded was 4 (> 75% sperm with progressive movement) and the mean motility duration was 40 ± 2 seconds with no significant differences among individuals between weeks.

**Figure 9.**
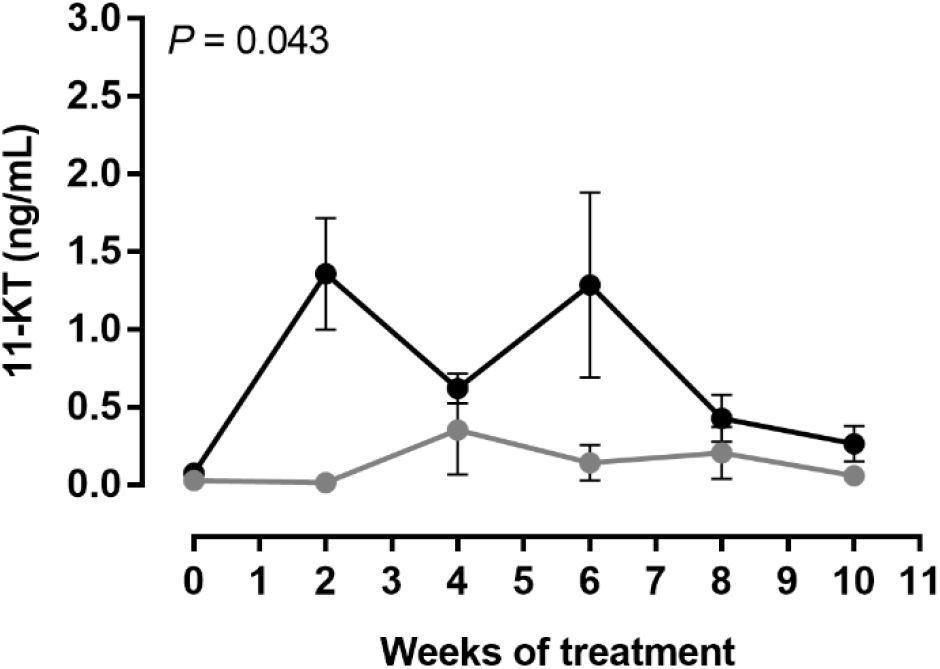
Mean (± SEM) plasma 11-KT levels of rFsh-treated (15 μg kg−1) males grey mullets and controls (n = 3-4). Treated males received weekly injections of rFsh (15 μg kg−1) and control males of CHO conditioned culture medium (1 mL fish-1). There are significant differences among treatments (two-way repeated measures ANOVA, P = 0.043, α = 0.05, statistical power = 0.66).

In Experiment 2, along the course of the treatment, all treated males (n = 4) produced sperm, which also coincided with an initial significate increase in 11-KT levels in the treated group (P = 0.006, α = 0.05, statistical power = 0.97) (before treatment: 2.2 ± 0.8; after 4 weeks: 10.5 ± 2.2 ng mL−1) in comparison with the control group (before treatment: 0.7 ± 0.3; after 4 weeks: 0.5 ± 0.2 ng mL−1). Sperm could be obtained by abdominal pressure in treated males after 3 weeks of treatment (50% of males), 4 weeks (75 %) and from the fifth week to the end of the treatment (100 %). From the third week of treatment to the fifth, first traces of sperm (mean 30.3 ± 12.3 μL) were highly viscous with a significantly higher concentration of spermatozoa (mean 2.1 ± 0.2 1011 spz mL−1) and a motility score of 2 to 4 (25 to > 75% motility). After six weeks, higher quantities of sperm where obtained (242.5 ± 70.9 μL) coinciding with a previous increase in rLh administration, which decreased (68.7 ± 13.7 μL) afterwards. Viscosity and spermatozoa concentration (2.3 ± 0.8 1010 spz mL−1) significantly decreased compared to the first weeks sperm was obtained. Motility score was 4 for all males until the end of the treatment. Mean duration of sperm motility was 89 ± 14 seconds during the 6 weeks that sperm was collected.

Assessment by CASA of the 10 samples collected from all four males with high motility score and ≥ 100 μL volume showed a mean motility percentage of 74 ± 0.01 %, VCL of 90.7 ± 3.3 μm s−1, VAP of 84.6 ± 5.5 μm s−1, VSL of 83.4 ± 6.9 μm s−1, LIN of 91 ± 0.5 %, WOB of 93.5 ± 0.1 % and STR of 97.9 ± 0.7 %.

### 3.5. *In vitro* fertilisation

The 0.5 mL aliquots of stripped eggs (1224 ± 150 eggs) were fertilised by mixing with 60 μL (20 μL/male) of pooled diluted stripped sperm (sperm 1:4 in Marine Freeze®) (3.8 ± 0.8 109 spz mL−1). The mean sperm to egg ratio at fertilisation was 189,521 ± 23,541 spermatozoa egg-1. After an incubation period of 22 - 23 hours (24°C), mean embryo survival rate was 0.4 ± 0.2 % (n = 3 females). At this age, the head region had formed and dark pigments covered almost all of the embryo and the oil globule (Fig 10A). Although, a single oil yolk globule was noticed in the majority of embryos, 28 % of the examined eggs presented multiple oil droplets. Mean fertilised egg diameter was 844 ± 4 μm. Hatching rate of the fertilized eggs, observed at 39 - 40 hours after fertilisation, was 70.8 ± 20 % (Fig. 10B). Mugil cephalus larvae at 1 dph had developing eye lens and a reduced yolk sac diameter (Fig 10C). At 2 dph the yolk and oil globule were still present, but mouth parts were completely formed with upper and lower jaws opened (Fig 10D). Survival rate of larvae decreased to 38.6 ± 22 % at 1 day post-hatching (dph) and continued decreasing to 4.1 ± 1.4 % (2 and 3 dph) until zero (4 dph).

**Figure 10.**
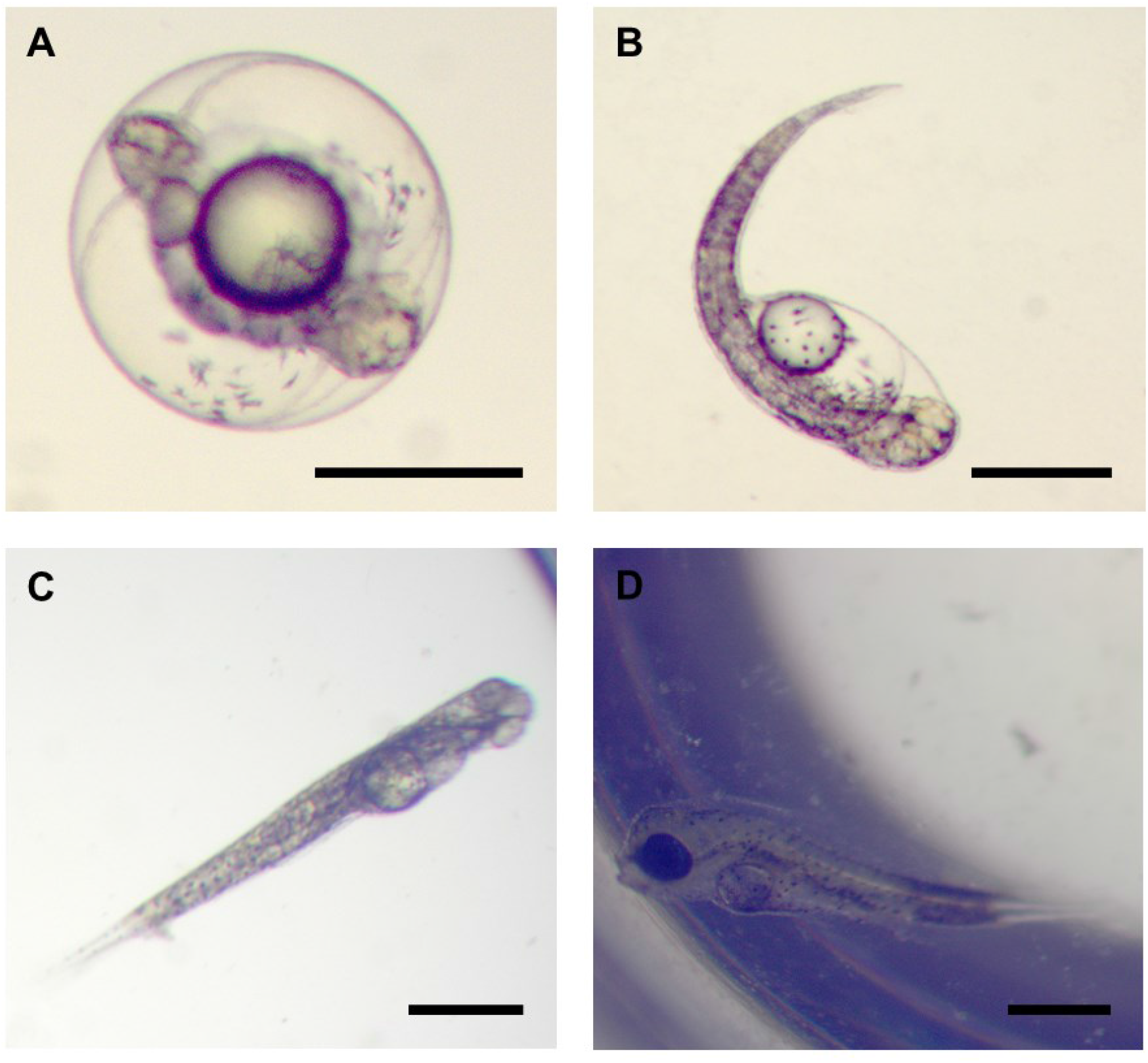
Developing *Mugil cephalus* embryos and larvae from Experiment 2. (A) Embryo at age 22 h post-fertilisation with head region formed and dark pigments covering almost all the embryo and on the oil globule. (B) Hatching at age of 40 hours post-fertilisation. (C) Larva after 1 dph. A decrease in yolk sac was observed and the eye lens formed. (D) Larva after 2 dph with well-developed eye, with mouth parts formed and opened. Oil globule was still present. Scale bar: 500 μm.

## 4. Discussion

The present study shows that rFsh drives oogenesis from early to late gonad developmental stages in female grey mullet, that rLh is influential to achieve oocyte maturation and ovulation and that rGtHs can be used to produce milt from male grey mullet. These findings are significant to both demonstrate the accepted roles of the Gths in teleost reproductive development and to provide tools for the control of reproduction in teleost species that experience reproductive dysfunctions early in the maturation process.

Flathead grey mullet is a species that exhibits severe reproductive dysfunctions in captivity that threatens the sustainability of mullet culture making it mostly dependant on wild captures. Despite of the present study being timed to coincide with the natural reproductive period, no reproductive development was observed in control females that remained arrested in primary growth or cortical alveoli stage and no sperm was obtained from control males when abdominal pressure was applied. All of the control fish had sufficient size, 35 - 49 cm for females and 32 - 42 cm SL for males, and condition to mature according to reported sizes of maturity; 27 - 35 cm standard length for females and 25 - 30 cm for males (Whitfield et al., 2012). The present study, encountered a more severe reproductive dysfunction than has been observed in other studies (Aizen et al., 2005). The severity of the reproductive dysfunction, highlights that in the present study, the application of rGths was critical in stimulating reproductive development in the experimental female fish and availability of sperm in males.

The hormone therapy to control the entire process of oogenesis was initiated with the application of rFsh. The administration of different doses of rFsh to examine the biological activity of this recombinant hormone, obtained a significant and prolonged (3 - 6 days) increase of E2 after injection of 12 - 15 μg kg−1. The increase in plasma E2 levels reflected the gonadotropic stimulation of the ovary by rFsh produced in the CHO system. The potent activity was further confirmed by the significant increase in E2 plasma levels in relation to the weekly administration of rFsh (15 μg kg−1 in Exp 1 and increasing doses 6, 9 and 12 μg kg−1 in Exp 2) to female grey mullets. The plasma levels of E2 seven days after rFsh application at week 4 were 272 ± 43 pg mL−1 in Exp 1 and 458.7 ± 113 pg mL−1 in Exp 2. In the present study, the rFsh-mediated increase of E2 plasma levels in females appeared to stimulate oocyte growth by the accumulation of lipid globules and yolk droplets, as E2 stimulates vitellogenin synthesis by the liver (Lubzens et al., 2010). The fact that the rFsh doses including lower rFsh doses in Exp 2 were sufficient to induce vitellogenesis may indicate that rFsh doses could be refined for future inductions.

In both experiments, oocytes grew from immature perinucleolar stage and/or cortical alveoli stage to advanced vitellogenic stages after rFsh administration. This oocyte growth was observed in eight (89%) of the nine treated females in both experiments. There was some variation in individual responses that ranged from a few more advanced females to two females (one in each experiment) that did not reach vitellogenic stages in the 4 - 5 week-period. Despite of this variation, the present study presents a considerable advance to successfully induce oogenesis in 89% of experimental fish with the application of rFsh in a teleost. The biological activity of rFsh applied to females of other fish species has been previously studied, but most studies have focused on in vitro approaches for receptor-binding capacity (So et al., 2005) and steroidogenic potency (Kasuto and Levavi-Sivan, 2005; Meri et al., 2000; Zmora et al., 2007) or in vivo short-term effects (Kazeto et al., 2008; Ko et al., 2007; Kobayashi et al., 2006, 2003; Molés et al., 2011) rather than in vivo long-term effects on gonadal development. When rFsh produced in other heterologous systems than CHO cells were tested in long-term treatments in different fish species, more time, dose and/or number of administrations were required to reach a less advanced stage of ovary development than in the present study. For instance, after 60 days of treatment with injections at 10-day intervals of rFsh produced in P. pastoris (10 - 20 μg kg−1) immature yellowtail kingfish oocytes developed to cortical alveoli stage (Sanchís-Benlloch et al., 2017). Weekly injections for 8 weeks at 100 μg kg-1 to juvenile grouper (Epinephelus fuscoguttatus) also induced development to the cortical alveoli stage (Palma et al., 2018). Recombinant Fsh produced in Drosophila S2 cell line (100 μg kg−1) induced early vitellogenesis in the Japanese eel after 56 days of treatment with a weekly administration (Kazeto et al., 2008) and rFsh (500 μg/kg) produced in HEK293 cells induced initial oil droplet stage in previtellogenic yellow shortfinned eels (Anguilla australis) after three weeks (Tuan Nguyen et al., 2020). These comparisons between the present study and other studies suggest a higher biological potency of rFsh produced in CHO cell lines as previously reported in some species (Molés et al., 2011).

Nevertheless, the administration of only rFsh in Exp 1 failed to complete oocyte growth as although oocytes developed until mid to late secondary growth, the cells appeared to be arrested in this stage and subsequently, a substantial number of atretic cells were found in the later weeks (weeks 9 - 11). These results agree with previously described E2 roles that did not include OM (Lubzens et al., 2010), but differ from those obtained by Das et al. (2014) who induced OM in M. cephalus post-vitellogenic oocytes that were incubated in vitro with E2. The fact that completion of oocyte growth could not be achieved using only rFsh suggested that, as previously described, OM and ovulation are Lh-dependent (Lubzens et al., 2010; Nagahama and Yamashita, 2008). According to Nagahama and Yamashita (2008), secretion of Lh from the pituitary coincides with a switch in the gonad steroidogenic pathway from the production of predominantly E2 during vitellogenesis to the production of progestin-like steroids, the maturation-inducing steroids (MIS). The MIS bind to oocyte membrane-specific receptors to activate the maturation promoting factor (MPF) that induces germinal vesicle breakdown and OM (Lubzens et al., 2010). Therefore, in Stage 2 of Exp 1 and Exp 2, we focused on the use of exogenous sources of Lh receptor agonists or hormones that may trigger the release of Lh from mullet pituitary with the aim to complete oocyte growth and induce OM.

The application of hormone treatments (GnRHa+MET or hCG) in Stage 2 of Experiment 1 failed to induce oocyte growth and OM. The oocytes remained arrested in the secondary growth stage of development with mean oocyte diameters of 425 ± 19 μm and an increasing incidence of atresia. It appeared that the developmental stage of the oocytes was not sufficient to respond to the hormone treatments that have been successful in a wide range of species that were arrested at a later developmental stage close to OM (Malison et al., 1998; Mañanós et al., 2009). Other studies on Mugil cephalus have recommended an oocyte diameter > 550 μm before OM and ovulation induction (Aizen et al., 2005; El-Gharabawy and Assem, 2006; Vazirzadeh and Ezhdehakoshpour, 2014). However, a wide range of other possible contributing factors can be cited, such as pituitary Lh content may have been low, the follicles were not receptive at the time of hormone application and did not stimulate the switch in gonad steroidogenic pathway to MIS or that the administration of rFsh complicated the switch as agonists of the Lh receptor also stimulated the Fsh receptor (Chauvigné et al., 2012; So et al., 2005). However, considering that the hormone treatments used in Stage 2 of Experiment 1 were applied to few fish, no conclusion can be drawn other than oocyte development was arrested with the application of only rFsh and no further development was observed.

In contrast, in Experiment 2, the co-administration of rLh with rFsh at advanced stages of vitellogenesis induced the completion of oocyte growth to a mean size of 609 ± 5 μm in eight (89%) of the nine females treated. Experiment 2 compared to Exp 1, appeared to show that the addition of rLh was required to increase maximum oocyte diameter to a diameter (>550 μm) that represents the completion of oocyte growth and a diameter from which OM has been observed to progress (Tamaru et al., 1993; Yousefian et al., 2009, present study). The increase in oocyte diameter and advance in development obtained in experiment 2, indicated that the completion of vitellogenic growth was dependent on Lh, which has not been previously described. The dosage and the time interval of rLh treatment applied to induce OM were based on previous studies (Chauvigné et al., 2017). However, since the half-life of rLh in plasma has not been determined in grey mullet, the most efficient hormone treatment (dose and timing) remain to be established. In relation to the induction of OM and ovulation, the rationale behind the treatment of rLh plus progesterone, a precursor of maturation-inducing steroids, was to induce the Lh-mediated up-regulation of genes associated with these processes and to avoid potential substrate-limiting factors for MIS synthesis. In Exp 2, only the five fish receiving the highest rLh dose (30 μg kg−1) with P proceeded to OM compared to three fish that received a lower rLh dose (15 μg kg−1) with P that did not develop to OM. This indicated that rLh dosage has a relevant effect and high doses were required. Recombinant Lh has been previously successfully used to induce OM and ovulation in bitterling (Rhodeus ocellatus ocellatus) (Kobayashi et al., 2006), common carp (Cyprinus carpio) (Aizen et al., 2017) and Malaysia catfish (Hemibagrus nemurus) (Salwany et al., 2019). However, the present study cannot confirm if a unique injection of rLh could have completed OM and ovulation without the need of progesterone application. Further work is required to fully understand the roles and administration of rFsh, rLh and P to successfully execute the steroid switch to induce OM and ovulation. The mean fecundity of the four (44%) females that were successfully induced with rLh and P to complete OM and ovulation was 742,900 ± 71,840 eggs kg−1 bw (~ 855,800 eggs female-1), which was within the range previously reported for M. cephalus, from 500,000 to 3,000,000 eggs female-1, that shows variation in relation to fish size and the technical procedures employed for egg collection (González-Castro and Minos, 2016).

The present study also provides evidence that Fsh is the major hormone to initiate vitellogenesis in teleosts. To date, no study has demonstrated that the exogenous application of just Fsh promotes the initiation of vitellogenesis and development through to late vitellogenic stages and that development progressed correctly to provide oocytes for the formation of viable eggs and larvae. The central role of Fsh in fish vitellogenesis is accepted (Mañanós et al., 2008; Lubzens et al., 2010) based on parallels drawn with other taxa, the synchronised increase in plasma Fsh and oocyte development found in many fish species, genomic approaches such as gene knockout to define Gths pathways (Zhang et al., 2015) and that rFsh induced partial development of vitellogenesis (Kazeto et al., 2008; Palma et al., 2018; Sanchís-Benlloch et al., 2017; Tuan Nguyen et al., 2020). However, some criticisms can be made as, many differences in the control in reproduction exist between taxa, synchronised increases in Fsh and oocyte development do not necessarily indicate cause – effect, vitellogenesis although delayed proceeded when the Fshb gene was knocked out to make Fsh-deficient zebrafish (Danio rerio) (Zhang et al., 2015) and previous studies did not induce the entire process of vitellogenesis (Kazeto et al., 2008; Palma et al., 2018; Sanchís-Benlloch et al., 2017; Tuan Nguyen et al., 2020). Therefore, the present study has added clear evidence to demonstrate the accepted function of Fsh by reporting in a teleost species that rFsh successfully induced the process of vitellogenesis from previtellogenic stages to advanced stages from which fertilised eggs and larvae were obtained.

Regarding males, the rGths treatments induced the production of milt for fertilisation procedures. The biological effects of rGths were evaluated through plasma 11-KT levels and by the presence of milt after abdominal pressure. The rFsh treatment in Exp 1 and rFsh with rLh in Exp 2, both significantly increased the levels of 11-KT, which is the major androgen responsible for testicular development (Aizen et al., 2005; Chauvigné et al., 2012; Mañanós et al., 2009; Schulz et al., 2010). In comparison, no sperm could be obtained from males in control groups. Other studies have induced or increased the production of milt obtained by abdominal pressure in immature Japanese (Hayakawa et al., 2008; Kamei et al., 2006; Kobayashi et al., 2010) and European eel (Peñaranda et al., 2018) and mature Senegalese sole (Chauvigné et al., 2018, 2017) after gonadotropin administration. The administration of rFsh alone induced the production of low milt volumes, whilst the additional administration of rLh increased milt volumes and decreased spermatozoa concentration probably due to a stimulation of the production of seminal fluid. The induction of spermiation by rFsh alone has also been demonstrated in the European eel (Peñaranda et al., 2018) and similarly the addition of rLh increased volumes and decreased spermatozoa concentration. The little seminal fluid produced in the present experiments could explain the higher sperm concentrations observed (in the range of 1010 and 1011 spz mL−1) with respect to that previously reported for this species

(108) (Ramachandran and Natesan, 2016). Nevertheless, the rGth treatments provided sperm for fertilisation procedures even though the number of males in the study was low. Curiously, the present study also indicated that there may be a sex specific contrast in the effect of rFsh, as in males rFsh alone induced the production of mature spermatozoa compared to females in which rFsh alone did not induce mature gamete production, and ovaries were arrested in late vitellogenesis and atresia was observed. However, further studies are required to examine and determine the existence of this sex specific difference and clarify the interactions amongst rGth levels and receptors or the mechanisms that may be responsible.

After hand stripping gametes (3 females and 3 males) and in vitro fertilisation, 0.4% of eggs developed embryos. The low percentage of eggs developing an embryo may be related to in vitro fertilisation procedures. The morphological aspect of the eggs appeared normal with the exception that 28% of the eggs had multiple oil droplets. In grey mullet, the manual pressure of artificial stripping increased the frequency of multiple oil droplets (Kuo et al., 1973) and multiple oil droplets were related with low egg survival (Nash and Shehadeh, 1980). Another aspect related to bad egg quality and in vitro fertilisation procedures is overripening (Ramos-Júdez et al., 2019). After ovulation, there is a period of egg ripeness with optimal viability after which the eggs overripen, losing quality and viability. This period of optimal egg quality for stripping varies among species, with temperature, between different stocks, holding conditions, hormone induction treatments and ideally should be defined for each situation (Ramos-Júdez et al., 2019). For example, latency to obtain good quality eggs can be as long as 5 - 15 days over a temperature range of 10 - 17 °C for rainbow trout (Oncorhynchus mykiss) (Samarin et al., 2008), 3 hours in meagre (Argyrosomus regius) at 18 °C (Ramos-Júdez et al., 2019) but only 30 min in white bass (Morone chrysops) at 22 °C (Mylonas et al., 1996). For the present treatment in M. cephalus, the timing of ovulation and optimal egg quality has not been previously defined. However, latency times have been reported for grey mullet using carp pituitary extracts with hCG or GnRHa (Karim et al., 2016), hCG (Kuo et al., 1973) and pituitary glands combined with synahorin and vitamin E (Liao, 1975) and times ranged from 30 to 48 hours after the initial priming dose and 12 to 26 h after the resolving dose. In the present study, eggs were stripped at 40 h and 48 ± 0.5 h from rLh administration (16 h and 24 h from progesterone). The female stripped at 40 h was induced to ovulate with PG-F2α and the stripped eggs had vitellogenin and oil in the process of coalescing apparently not having completed maturation and hydration when the oocytes were ovulated. On the contrary, at 48 ± 0.5 h after the rLh injection, low fertilisation rates were obtained. This was at the limit of the period of good egg quality that has been found with other hormone treatments (30 to 48 hours), which may indicate that the 48-h stripping time was late and that the eggs were undergoing overripening. However, it cannot be discounted that the egg quality was low due to aspects of the rGth induction protocol. Therefore, further studies to determine the timing of ovulation and the window of good egg quality are crucial to determine the quality of eggs that can be achieved with rGth based therapies.

Fertilised grey mullet egg diameter has been reported to vary from 0.65 - 1.08 mm differing with different geographical areas (González-Castro and Minos, 2016). In the present study, the fertilised eggs ranged in diameter from 0.82 to 0.88 mm at a temperature of 24°C and salinity of 36 ‰. Hatching was 39 - 40 hours after fertilisation at 24 °C, which is in agreement with previous reports of hatching time: 34 - 38 h at 22 - 24.5 °C and 49 - 54 h at 22.5 - 23.7 °C (González-Castro and Minos, 2016). High mortalities were found at two and three-days post hatching (dph), which coincides with the period that mouth, upper and lower jaws opened although the yolk sac was still present. These high mortalities were probably due to starvation as no food was offered and survival depends on the availability of external food organisms to larvae on the second-day, 36 hours post-hatch, before the completion of yolk sac absorption (Abraham et al., 1999).

In conclusion, the present study reports that treatment with rGths was able to induce full oogenesis to produce viable eggs and larvae in a teleost. The control of the entire reproductive process using rGths, and particularly the initiation of vitellogenesis, development through to late stages with rFsh and the completion of oocyte growth with rLh offer further data about the roles of the Gths in teleost oogenesis. A refined protocol based on the present study could provide full reproductive control of flathead grey mullet held in intensive aquaculture facilities. In addition, these findings raise the possibility of using the rGth treatments for species that present similar reproductive disorders in aquaculture, the aquarium industry and for the conservation of endangered species.

## Acknowledgments

The authors would like to thank Josep Lluis Celades and the IRTA staff for technical help. Special thanks to Narciso Mazuelos from Finca Veta La Palma (Isla Mayor, Sevilla) for providing us with mullet broodstock. This study has been supported with funding from the European Union’s Seventh Framework Programme for research, technological development and demonstration (GA 603121, DIVERSIFY to H.R. and N.D.), the Spanish National Institute for Agronomic Research (Instituto Nacional de Investigación y Tecnología Agraria y Alimentación - INIA)-European Fund for Economic and Regional Development (FEDER) (RTA2014-0048 to N.D.), the Spanish Ministry of Science and Innovation (Ministerio de Asuntos Economicos y Transformación Digital - MINECO) (RTI2018-094710-R-I00 to N.D.) and Rara Avis Biotec S. L. (I.G., Valencia, Spain). Participation of S.R. and W.G. was funded, respectively, by a PhD grant from AGAUR (Government of Catalonia) co-financed by the European Social Fund and a PhD grant from the National Board of Science and Technology (CONACYT, Mexico).

